# Disulfide reduction allosterically destabilizes the β-ladder sub-domain assembly within the NS1 dimer of ZIKV

**DOI:** 10.1101/2020.05.01.072405

**Authors:** P Roy, S Roy, N Sengupta

**Affiliations:** IISER Kolkata, India; IISER Kolkata; Centre for Excellence in Basic Sciences, Mumbai

## Abstract

The Zika virus (ZIKV) was responsible for a recent debilitating epidemic that till date has no cure. A potential way to reduce ZIKV virulence is to limit the action of the non-structural proteins involved in its viral replication. One such protein, NS1, encoded as a monomer by the viral genome, plays a major role *via* symmetric oligomerization. We examine the homodimeric structure of the dominant β-ladder segment of NS1 with extensive all atom molecular dynamics. We find it stably bounded by two spatially separated interaction clusters (C1 and C2) with significant differences in the nature of their interactions. Four pairs of distal, intra-monomeric disulfide bonds are found to be coupled to the stability, local structure, and wettability of the interfacial region. Symmetric reduction of the intra-monomeric disulfides triggers marked dynamical heterogeneity, interfacial wettability and asymmetric salt bridging propensity. Harnessing the model-free Lipari-Szabo based formalism for estimation of conformational entropy (S_conf_), we find clear signatures of heterogeneity in the monomeric conformational entropies. The observed asymmetry, very small in the unperturbed state, expands significantly in the reduced states. This allosteric effect is most noticeable in the electrostatically bound C2 cluster that underlies the greatest stability in the unperturbed state. Allosteric induction of conformational and thermodynamic asymmetry is expected to affect the pathways leading to symmetric higher ordered oligomerization, and thereby affect crucial replication pathways.

**Statement of significance:** Controlling viral pathogenesis remains a challenge in the face of modern-day epidemics. Though cumbersome and fraught with misleads, most therapeutic endeavors lean towards the design of drug molecules targeting specific proteins involved in viral pathogenesis. This work demonstrates an alternative approach, namely the usage of allosteric intervention to disrupt the binding integrity of the primary domain of the non-structural NS1 protein dimer crucially important in ZIKV virulence. The intervention, triggered by symmetric reduction of the internal monomeric disulfide bonds, results in weakening and distortion of the distal binding interfaces. It further introduces marked structural and entropic asymmetry within the homooligomeric unit, precluding the formation of higher ordered oligomers of high symmetry. The results have important ramifications for consolidated efforts at limiting ZIKV virulence.

## Introduction

Protein oligomerization is a common evolutionary process that underlies biological functionality, and in particular situations, potentially triggers disease onset with production of non-functional aggregates. In a typical cell, more than 35% of the proteins may oligomerize and attain stability against denaturation (1). Functional oligomeric states can vary from small oligomers to higher ordered assemblies such as stable amyloid fibrils and viral capsids (2,3). Homooligomeric assemblies are involved in diverse cellular functions including enzymatic activity; molecular recognition; formation of transmembrane channels and carriers; and in signaling pathways (4–6). The nature of oligomerization depends on the contributing polypeptide sequence, and the resultant state is stabilized by superposing non-covalent interaction networks that may extend from the oligomeric interface to interior regions via allosteric networks. The association of sequentially identical monomers into structurally stable, symmetric units has been attributed to finite control of assembly (1,7).

Small genome sized infectious agents such as viruses are known to exploit symmetric supramolecular organization as an evolutionarily advantageous strategy (8). Such a mechanism has been associated with the Zika virus (ZIKV), an arbovirus within the flavivirus family that triggered a recent epidemic (9). The second ZIKV outbreak gave rise to the Guillen-Barre (GB) syndrome with devastating neurological consequences (10). The persisting virulence of Zika has triggered urgent preventive and remedial efforts towards restricting infection (11). After entering a host cell, the 11 kb genome of this flavivirus synthesizes three structural (envelop, capsid and pre-membrane) and seven non-structural (NS1, NS2A, NS2B, NS3, NS4A, NS4B, and NS5) proteins (12). The structural proteins take part in virion formation, whereas the non-structural proteins are primarily involved in replication and orchestration of viral morphogenesis (13).

Recent mutational and phylogenetic studies indicate that ZIKV spread is facilitated by the non-structural protein NS1, whose mutations can enhance antigenemia and inhibit the interferon beta-induction (14,15). NS1 is a multifunctional, yet highly conserved protein within the flavivirus family (16). Its general roles include immune evasion and invasion, formation of viral replication complex, and interactions with ribosomal subunits. NS1 spontaneously form multimeric structures that are capable of localization in various cellular regions, and form trimers of dimers (hexamers) in lipid rich extracellular milieu; the latter is a potential biomarker in diagnosis. Disturbing NS1 oligomerization could, therefore, provide crucial therapeutic leverage towards halting flaviviral infection, particularly with ZIKV.

The mature form of NS1 presents a dimer of identical monomeric subunits (17). The structurally conserved monomer has three domains: a small N-terminal β hairpin (residues 1-30); a wing domain (residues 37-175), and the largest, C-terminal β-ladder domain (residues 180-352). The domains are connected through first (residue 31-36) and second (residue 176-180) connector subdomains (18). The β-ladder domain is the dominant dimerization feature within the hexamer, presenting the vast majority of contact points in the dimeric interface (18–20). Weakened dimerization is brought about by β-ladder mutations such as by insertion of the M2e peptide of influenza A virus (21). Single point mutations within NS1 can ablate dimer formation and reduce virulence (22). Structurally, this domain consists of 10 β strands arranged like a ladder rung and connected through short loops or turns except a 53 residue long (179 to 223 residues) loop called as “spaghetti loop” between β13 and β14 strands. The presence of the spaghetti loop gives rise to two different surfaces: the membrane side of the continuous β sheet and the luminal side of irregular loop surface (18).

It is noteworthy that four of the six internal disulfides in the NS1 monomer are located in the β ladder domain (18,20). Disulfides play significant roles in protein folding and stability (23,24). Their post-translational formation increases the conformational barrier for unfolding at the expense of entropic penalty, driving polypeptide chains to gain functionally stable structures (25,26). Mutagenesis studies of cysteine residues in the highly homologous dengue NS1 reveal that the β-ladder disulfides crucially influence NS1 maturation and oligomerization (27). To our knowledge, however, insights underlying the inherent stability of the dimeric interface of ZIKV NS1 is lacking.

Herein, we report our detailed molecular dynamics simulation studies unearthing the effect of disulfide reduction on structural integrity of the ZIKV β-ladder domain. We first unearth the key interactions that stabilize the interface, followed by analyses of the effects of symmetric disulfide reductions. We conclusively demonstrate that distal disulfide reductions are capable of allosteric destabilization of the dominant electrostatic cluster at the interface. Further, the allosteric effects are associated with a breakage in structural and entropic symmetry between the monomeric units. This is a first report uncovering and leveraging hidden allosteric interactions to destabilize a crucial element of ZIKV activation, and holds potential for control of downstream virulence.

## Methods

### I. System setup and MD Simulations

Starting coordinates for the β-ladder domain were taken from the experimentally resolved crystallographic structure of the ZIKV NS1 dimer (PDB code: 5GS6) from Protein Data Bank (18). Residues numbered 176 to 352 that comprise the β-ladder domain were taken from both monomers without the glycan linkage at position 207. The β-ladder dimer was solvated in an orthorhombic box with TIP3P water molecules (28). A minimum distance of 15 Å between any protein atom and the box edge was ensured, and 8 Na^+^ ions were added to neutralize the system. The box size at setup measured 118×76×89 Å^3^. The same procedure was followed to generate each system. Disulfide residues in each of the ladder domain are in Cys179-Cys223 (DS1), Cys280-Cys329 (DS2), Cys291-Cys312 (DS3) and Cys313-Cys316 (DS4). These were reduced by adding hydrogens to the sulfur atoms to yield the DS1^R^, DS2^R^, DS3^R^, and DS4^R^ systems, respectively; the original system was labeled ‘WT’ (or wild type). The system preparation and visualization were done using VMD (29). The two monomers are designated M1 and M2. The NAMD 2.12 simulation package (30) and CHARMM 22 with CMAP (31,32) correction force field were used for the MD simulations. Each system was energy minimized for 30000 steps with the conjugate gradient method and equilibrated at 310 K at a pressure of 1 atm for 20 ns. Each production run then proceeded in the isothermal-isobaric (NPT) ensemble for 600 ns. Langevin Dynamics was used to maintain a constant temperature of 310 K with a collision frequency of 1 ps^−1^. A constant pressure of 1 bar was applied by the Nosé-Hoover Langevin piston method (33). A 2 fs of time step was used and the generated trajectories saved at 1 ps intervals. Orthorhombic periodic boundary conditions were applied. Non-bonded interactions were calculated within a 12.5 Å cutoff distance and smoothing started at 10.5 Å. Long-range electrostatic interactions were considered with the Particle mesh Ewald (PME) summation method (34). The SHAKE algorithm was used to constrain bonds involving hydrogen atoms (35). Two independent production runs amounting to a cumulative 1.2 μs simulations were generated for each system.

### II. Analyses

#### Differential RMSD

Conformational fluctuation between a reference and an evolving protein structure is usually quantified with the backbone root mean squared deviation (RMSD). To quantify conformational variation between two co-evolving monomers in a homooligomer trajectory, we used a modified form termed differential RMSD (d_RMSD_) evaluated at each time point ‘*t*’ as,

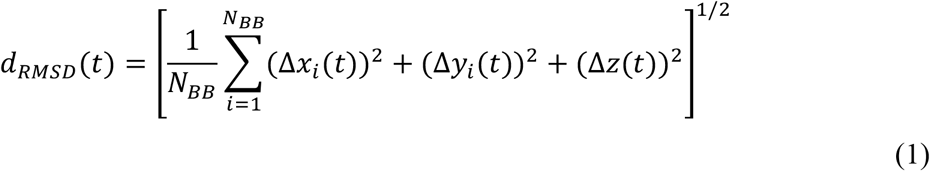

Here, *Δx_i_(t*), etc., represent the difference in Cartesian coordinates between the *i^th^* backbone heavy atom at a time ‘*t*’ after one monomer has been reoriented and superposed on the other in the least squares fit sense. *N_BB_* is the number of backbone heavy atoms.

#### Secondary structure persistence

The structural persistence of a protein over a reference (experimental) structure is defined as (36),

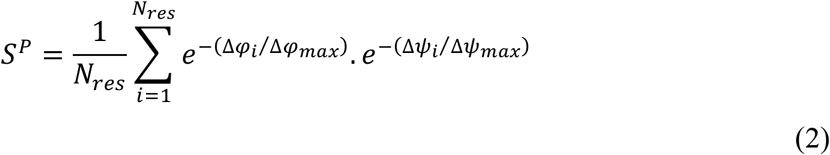

Here, *N_res_* represents the total number of residues; (Δ*φ_i_*, Δ*ψ_i_*) represent the magnitudes of changes in backbone dihedral angles of residue *i* over the reference; and (Δ*φ_max_*, Δ*ψ_max_*) represents the maximum allowed changes of the torsional angles in the Ramachandran space ignoring the direction of rotation. Therefore, an *S^P^* value of 1.0 represents a conformational state identical to the reference; lower values represent decreased overall persistence (37).

#### Salt bridge occurrence

A sidechain mediated hydrogen bond was considered a salt bridge when two oppositely charged residues come in contact within a donor-acceptor distance of 3.2 Å and a donor-hydrogen-acceptor angle not more than 150°. The salt bridge utility within VMD was used. A value of ‘1’ is assigned if a residue pair satisfies the salt bridging criteria and ‘0’ otherwise.

#### Essential dynamics

Principal component analysis (PCA) was used to identify independent modes of collective motion and thereby the essential dynamics. Briefly, the variance-covariance matrix of the backbone C_α_ atom fluctuations is first diagonalized and its eigenvalues and eigenvectors determined. The eigenvector corresponding to the largest eigenvalue is considered as the first principle component (PC1). The two extreme projections sampled along PC1 represent the bounds of the motional fluctuations and are graphically depicted as porcupine plots using PyMol. The porcupine plots are superimposed on the original structures to depict the extent of the deviation. The GROMACS (39) tools gmx_covar and gmx_anaeig were used in this analysis

#### Solvent accessibility and interfacial water molecules

The surface area of a protein region exposed to solvent, or the solvent accessible surface area (SASA) is estimated with VMD using a spherical probe of radius 1.4 Å. The water molecules at binding interfaces are identified as common waters shared by the participating residues. For the undistorted interfaces (WT and DS4^R^), common water molecules within a cutoff distance 2.5 Å of the residues were considered. For systems where the original interfacial contacts were substantially lost, shared water molecules within a cutoff distance of 3.0 Å were considered.

#### Conformational entropy

Conformational entropy (S_conf_), a key thermodynamic parameter governing protein function, can be challenging to estimate with conventional methods. Herein, we exploit a novel approach that recognizes the correlation between NMR estimates of local protein dynamics and S_conf_ (40). This model-free method can be used in conjunction with MD simulation data; benchmarked studies report excellent agreement with experiments (41,42). Briefly, the method utilizes the Lipari-Szabo squared generalized order parameter (*O^2^*) representative of the local internal motions of a protein (43). Following the removal of overall translation and rotation of the protein molecule, the order parameter (*O_i_^2^*) for an *i^th^* residue was estimated from an ensemble of the Cartesian projections of a bond vector. The generalized order parameter signifies the range of structural states that can be explored by capturing the motion of the interaction vector within the approximation of “diffusion in a cone”; its value varies from 0 (complete isotropic disorder) to 1 (complete rigidity) within the molecular frames of interest. Averages, indicated with angular brackets, were done over blocks of 10 ns of the equilibrated simulation data.

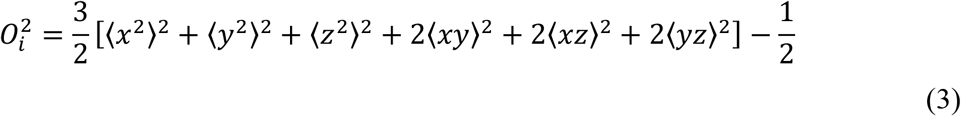

The total backbone conformational entropy (S^BB^), based on projections of the NH bond vector of *N* residues, is then calculated as,

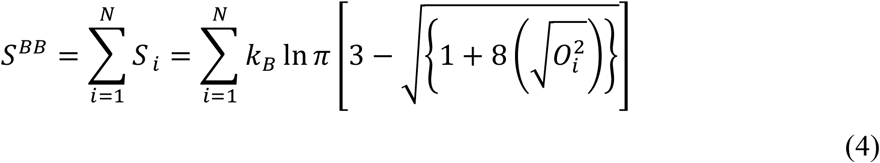

Here, *k_B_* is the Boltzmann’s constant. The total sidechain entropy is estimated by invoking the recent ‘entropy meter’ formalism that hinges on sidechain end methyl groups capturing the essence of local internal motion and is based on the CH bond projections.

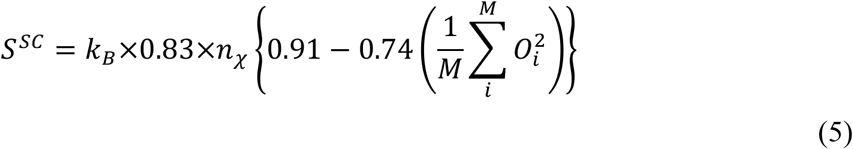

Here, *M* denotes residues of methyl bearing sidechain; *n_χ_* represent the total number of *χ* angles of the M residues (41,42).

## Results and Discussion

### I. Interactions underlying stability of the ladder dimer

We begin by examining the structural stability of the β-ladder dimer. As depicted in Fig. 1 *A*, the conformation obtained at the end of the simulations converge with the initial starting conformation. This is corroborated by the dimer’s low backbone root mean squared deviations (RMSD) and the high value of the overall structural persistence (S^P^); temporal evolution over a simulation is presented in Fig. 1 *B*. The mean inter-monomer interaction energy of this domain represents ca. 86% of the interaction strength of the full NS1 dimer (see Table S1 in the Supporting Material). These observations are consistent with previously mentioned experimental results indicating its significant contribution to NS1 stability (18).

**Figure 1.**
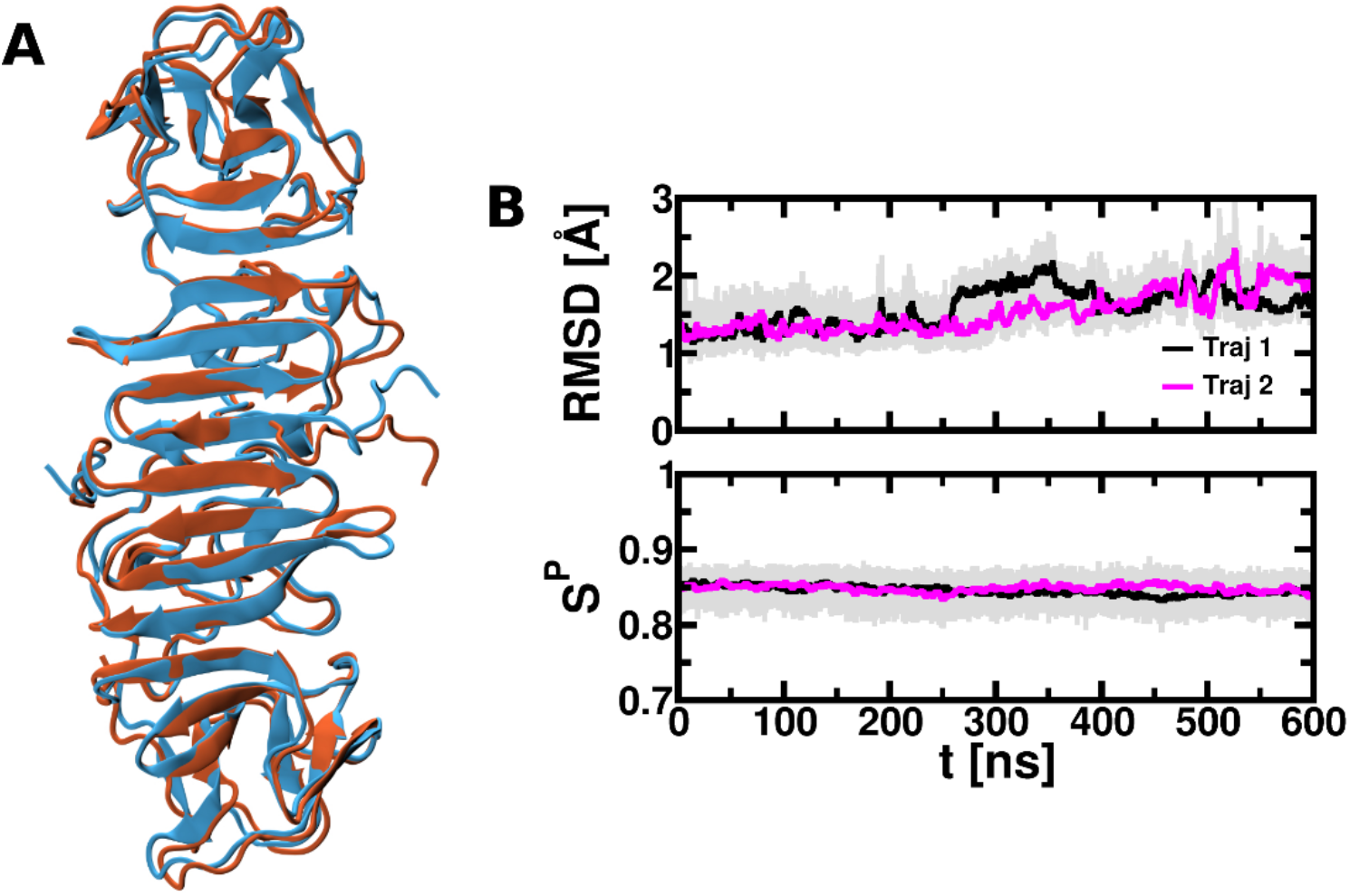
Structural stability of β-ladder dimer along the simulation trajectory. Superimposition of first (blue) and last (orange) conformations (*A*); time evolution of backbone RMSD (top panel; *B*) and structural persistence (bottom panel; *B*). The running averages done over 1 ns for each of the trajectories

In order to elicit the nature of interactions underlying NS1 stability, we investigated the inter-residue contacts between the two β-ladder monomers; two residues whose Cα atoms lie within 7 Å of each other are considered a contact pair. The contact probability map (Fig. 2 *A*) indicates the presence of two contiguous contact clusters, henceforth labeled C1 and C2, that can be distinguished in terms of their location, contact strengths, and nature of inter-residues interactions (see Fig. 2, *B-E*). The C1 cluster has 45 contact pairs and is localized at the interface region of the β face, whereas the C2 cluster has 29 contact pairs localized in the loop face characterized by the presence of a single 3_10_ helical turn (residue 227 to 229). However, only 27% of the pairs in C1 have a contact probability of 0.9; the corresponding value for C2 is 72%. We further note that in C1 and C2, the number of purely apolar contacts are 14 and 1 respectively, whereas the number of pairings between charged or polar residues is 7 and 21 respectively. This analysis indicates the predominantly electrostatic nature of the C2 cluster in comparison with C1.

**Figure 2.**
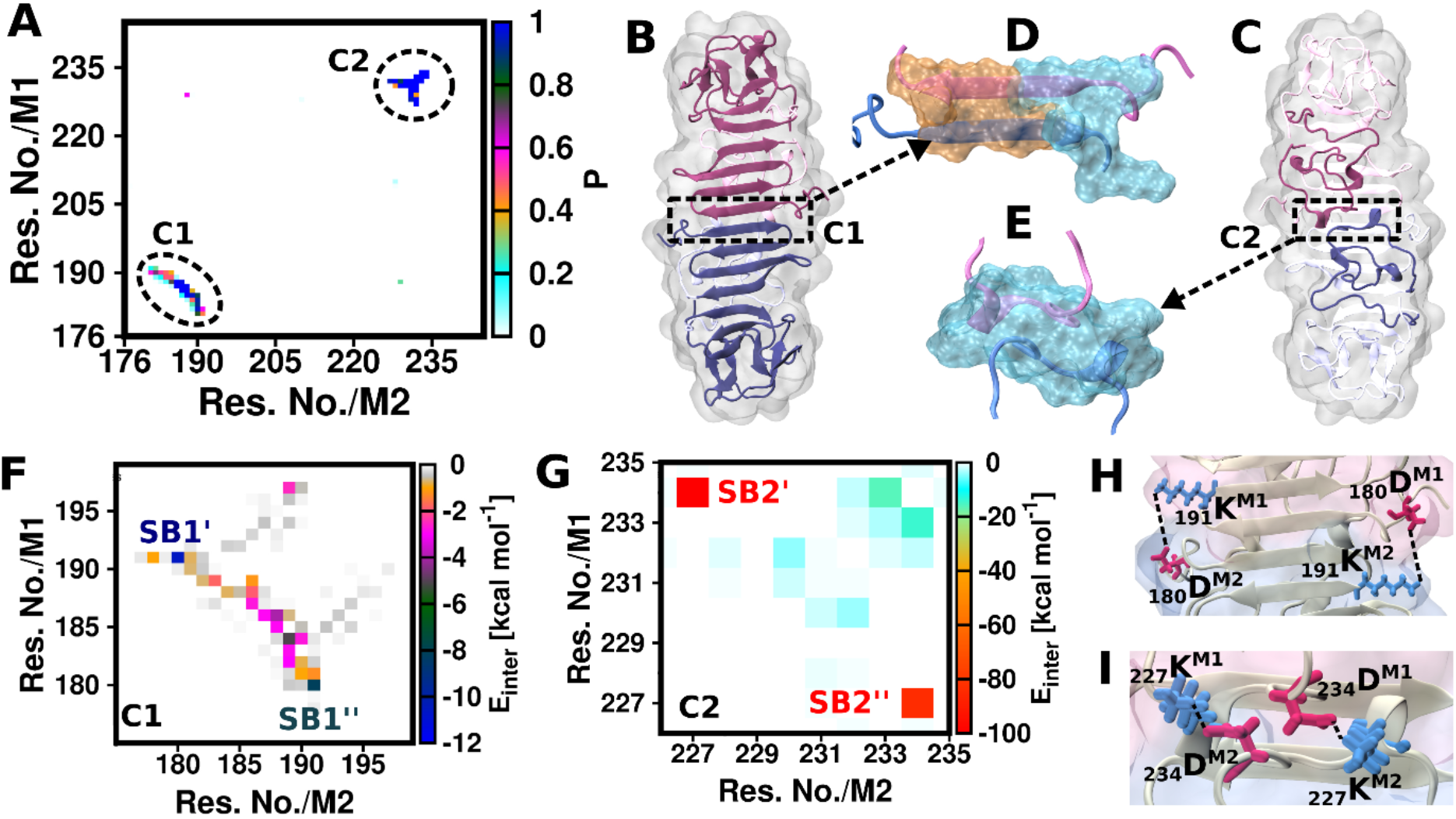
Binding hotspots of the β-ladder dimer. Pairwise inter-residue contact map between the monomers (*A*); location (*B* and *C*) and nature (*D* and *E*) of C1 and C2 contact cluster; pair-wise interaction energy contact map of C1 and C2 (*F* and *G*); representative snapshot depicting the location of key salt bridges of C1 and C2 (*H* and *I*). The system is shown in cartoon representation with distinct color of monomer 1 (M1, in mauve) and monomer 2 (M2, in blue); highly probable contacts (P>0.9) of C1 and C2 shown in surface where orange and cyan indicates hydrophobic and polar contact pair respectively. The positively and negatively charged residues colored as blue and red in the salt bridge rendition; hydrogen bonds are shown in dotted black line.

In Figs. *2F* and *G*, we depict the pairwise energetic contributions within each cluster estimated from the non-bonded energy terms of the interaction. In C1, nearly all the contacts contribute between −2 and −5 kcal mol^−1^, while a contribution of −12 kcal mol^−1^ is owed to a salt bridge, a polar contact (Lys191-Asp180) that lies at the cluster edge. In contrast, most residue pairs in C2 contribute an interaction of at least −20 kcal mol^−1^. C2 shows that it is characterized by two inter-monomer salt bridges (Lys227-Asp234) that contribute significantly to the interaction strength. Overall, the clusters represent the predominant binding sites of the β-ladder interface and contribute to ca. 98% of the total interaction energy; C1 and C2 contribute ca. 23% and 75%, respectively. Representative snapshots of the salt-bridge pairs of C1 and C2 are shown in Fig. 2, *H* and *I* respectively. The Lys227-Asp234 salt bridge is present in the 3_10_ helical turn of the C2 cluster. It is worthwhile noting that the structural integrity of the interfaces is not lost during the course of the simulations and the internal monomeric stability accompanies symmetric organization of contacts in C1 and C2.

### II. Disulfide reduction: evidence of long-range perturbation

The analyses thus far strongly suggest that perturbations to C1 and C2 could be key to disrupting NS1 dimeric integrity. While one may consider the development of small molecules that specifically bind to the clusters, targeted drug development can be cumbersome and fraught with misleads. In this light, we note recent endeavors towards allosteric control of protein dynamics and stability (44). Allostery is a thermodynamic phenomenon that can be modulated not only by small molecule and metal ion effectors but also by deploying altered solvent conditions, mutations, and selective covalent modifications (38,45–47). Resulting perturbations in dynamical coupling between key distal protein sites are accompanied by minimal changes to the protein’s internal structure. Allosteric modulation may be a more advantageous strategy over direct target of buried regions with small molecules.

Controlled allostery may be achieved by selective disulfide reduction within protein interaction networks (38,48). Disulfide contributes crucially to protein architectural stability and their cleavage can shift the thermodynamic balance towards non-native states. Interestingly, the highly homologous dengue NS1 shows reduced disulfide stability upon Cys to Ala mutations (27). We note that each β-ladder monomer of NS1 incorporates four internal disulfides, labeled henceforth as DS1 (Cys179-Cys223), DS2 (Cys280-Cys329), DS3 (Cys291-Cys312) and DS4 (Cys313-Cys316); see Fig. 3 *A*. The DS1 and C2 are positionally separated within the spaghetti loop region, whereas DS2-4 are present in the C-terminal region. In the β-ladder monomers, distributions of the S-S distance (*d_S-S_*) indicate high stability and low strain in all four bonds (Fig. S2); this is supported by DSE analysis (Table S2; see SI for more information). The mean center of mass distance between each disulfide and the binding hotspots C1 and C2 lies within 18-28 Å (Fig. S3). In the context of these observations, we systematically investigate the effect of symmetric DS1-4 reduction on the thermodynamic and structural stability of the β-ladder dimer.

**Figure 3.**
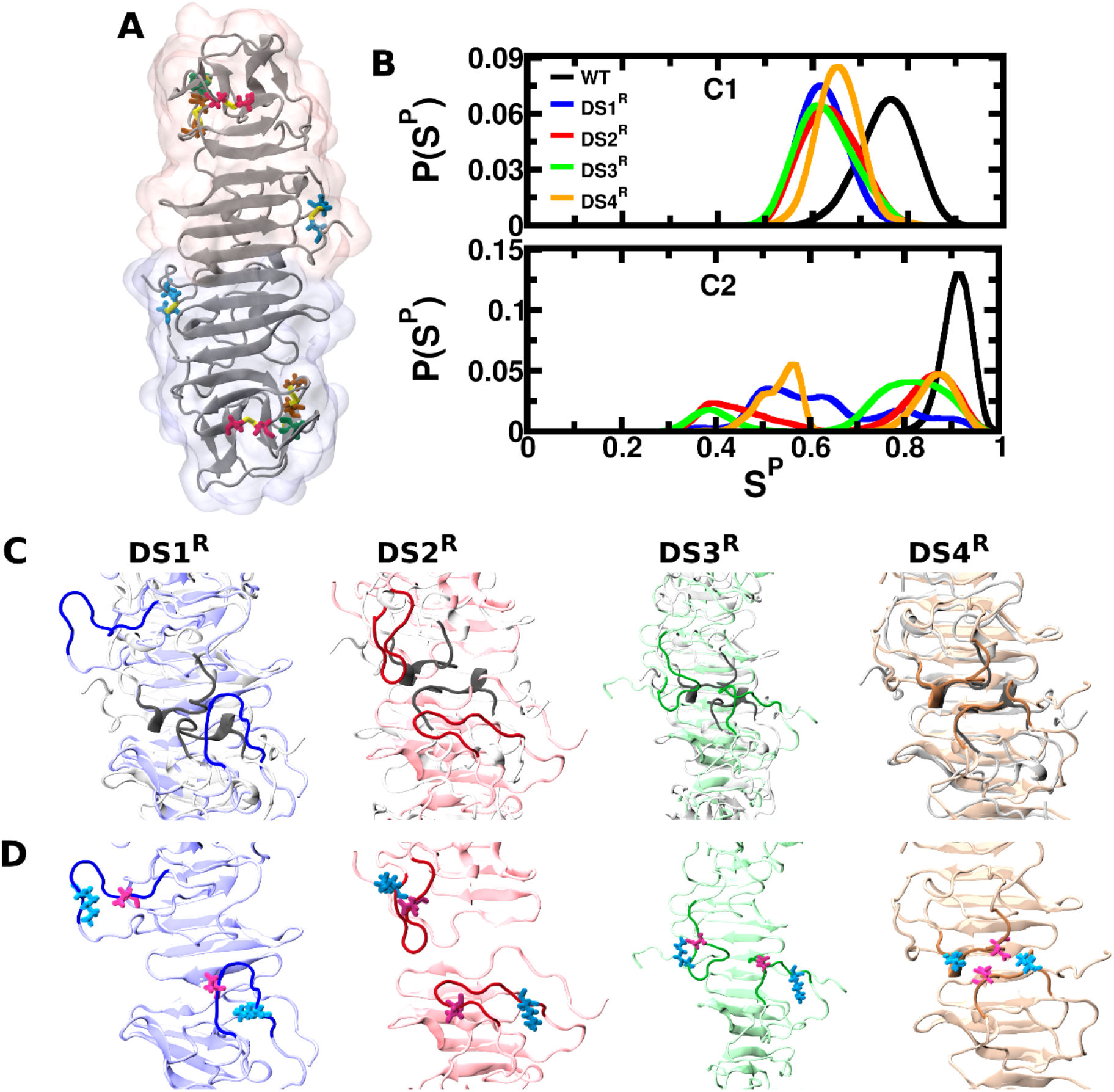
Effect of symmetric reduction of disulfides on the β-ladder dimer. Location of disulfides on the β-ladder dimer (*A*); Normalized distribution of structural persistence of C1 and C2 region (*B*); conformational plasticity of 3_10_ helix over the WT (*C*); and relative effect on the salt-bridge of C2 (*D*). Color code: DS1 (blue), DS2 (red), DS3 (green), DS4 (orange), Asp (magenta), Lys (blue2) and disulfide (yellow).

We first compared the change in the overall structural integrity of disulfide reduced (henceforth DS1^R^, DS2^R^, DS3^R^, and DS4^R^), and the unperturbed (WT) states; see Fig. S4. Although greater spread in the distributions of S^P^ indicates an increased conformational heterogeneity over WT, in a broad sense, disulfide reduction does not trigger major structural changes. Localized effects on the structured regions at the interfaces (β-strands in C1 and 310 helical turn in C2) are however more significant (Fig. 3 *B*), with sharp shifts from the WT and increased fluctuations. We observed greater susceptibility towards structural change in the C2 in DS1^R^. Inspection of the superposed first and last snapshots of each system indicates a noticeable loss in structure, particularly in the 3_10_ helical region of C2, for all reduced systems except DS4^R^ (Fig. 3, *C* and *D*). The intriguing local structural loss in key regions triggered by distal disulfide reduction lead us to investigate potential regulatory effects of the process. In Table 1, we compare the probabilities (P_50_) of 50% weakening of interaction strength after 100 ns, and the mean values obtained by combining the final 500 ns from each trajectory. While for the hydrophobic cluster C1, this probability is comparable for all systems, there are marked differences in the electrostatically stabilized C2 cluster. The P_50_ value for C2 is 0.02 and 0.09 in WT and DS4^R^ respectively but is 0.36, 0.47, and 0.55 for DS1^R^, DS2^R^, and DS3^R^, respectively.

**Table 1.** The probability of 50% weakening (P_50_) from of the initial interaction strength within 100 ns of simulations, and average interaction energy (<E_inter_>; in kcal mol^−1^) between the monomers of C1 and C2 region of β-ladder dimer calculated. Standard deviations of <E_inter_> are provided within braces.

| System | $P_{50}$ | | $\langle E_{\text{inter}} \rangle$ | |
| --- | --- | --- | --- | --- |
|  | C1 | C2 | C1 | C2 |
| WT | 0.28 | 0.02 | -59.47 (31.38) | -234.01 (53.76) |
| DS1<sup>R</sup> | 0.7 | 0.36 | -14.29 (25.45) | -156.29 (122.72) |
| DS2<sup>R</sup> | 0.4 | 0.47 | -22.75 (24.0) | -126.8 (113.43) |
| DS3<sup>R</sup> | 0.3 | 0.55 | -32.52 (26.08) | -68.86 (72.35) |
| DS4<sup>R</sup> | 0.17 | 0.09 | -40.55 (36.61) | -149.58 (65.04) |

The data thus far indicate that DS3^R^ has a relatively higher propensity of allosteric perturbation arising from disulfide reduction. For a clearer understanding, we evaluated the interaction maps of C1 and C2 of the reduced systems. The results presented in Fig. 4 are compared to the WT results in Fig. 2, *F* and *G*. Most major WT contacts in C1 are lost in DS1^R^-3^R^, with the emergence of new contacts in DS1^R^. While DS4^R^ retains most of the native C1 contacts, the Lys191-D180 salt bridge is faintly weakened. A more distinct alteration emerges in the C2 comparison. The contacts are nearly all lost in DS3^R^, and only transiently retained in DS1^R^ and DS2^R^. In contrast, DS4^R^ retains most native C2 contacts with high probability, although one of the two Lys227-Asp234 contacts are weakened. We note that the stability of the Lys227-Asp234 contacts in C2 are correlated with the structural persistence of the 3_10_ helical turn.

**Figure 4.**
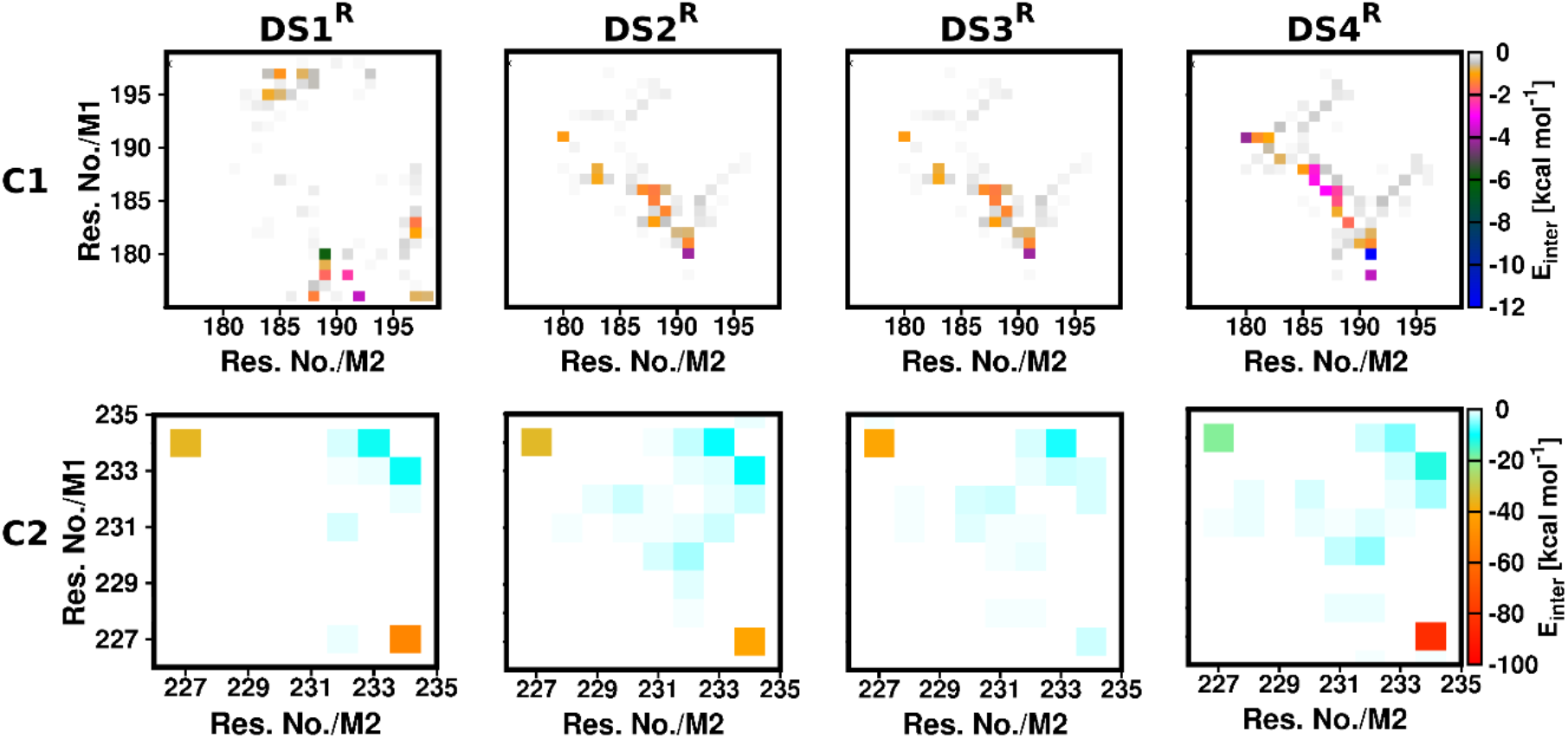
Contact map of pair-wise interaction energy for C1 (upper panel) and C2 (lower panel) of the disulfide reduced systems.

### III. Allosteric induction of homooligomeric asymmetry

With few exceptions, protein homooligomerization is concurrent with symmetric association and dynamics of non-covalent interactions (49). Structural symmetry in oligomers is generally associated with fast folding and attainment of the lowest energy state in a funneled energy landscape. Symmetry is further associated with cooperativity, stability against denaturation, and diminished solvent interactions (1,50). Homooligomerization appears to be prevalent in viral replication and pathogenesis (51,52). Intriguingly, it had been suggested by Watson and Crick that small viral genome is more conducive to attaining protein multi-functionality *via* protein homooligomerization than by direct translation of large protein units (50). It is further noted that conformational symmetry perturbation or breakage is harnessed in several biological processes including ligand binding, nonspecific DNA binding, enzyme activation, and signaling (53–57).

The interactions underlying the β-ladder dimer in the WT system display a high degree of symmetry, indicating its essence in higher ordered organization. It is noteworthy that distal disulfide reduction analyses so far suggest a propensity for symmetry loss of clusters C1 and C2 (see Fig. 4). For insights, we evaluated the differential RMSD (d_RMSD_) between the monomer pairs at each point along the simulation trajectories; see Fig S5. In Fig. 5 *A*, we compare distributions of d_RMSD_ values for all five systems; the median positions of DS1^R^-4^R^ are clearly at higher d_RMSD_ values over the WT. The mean d_RMSD_ values for WT are 1.53(±0.41) Å, and for DS1^R^-4^R^ are 4.91(±1.52) Å, 2.66(±0.65) Å, 3.05(±0.6) Å, and 2.41(±1.49) Å, respectively, indicating a clear increase in dynamical asymmetry between the monomers upon symmetric disulfide reduction (see Fig. S6 for cluster-wise dRMSD distribution). We further compared the essential dynamics of both clusters represented as porcupine plots by cumulatively combining equal segment of data from latter parts of both trajectories; this is presented in Fig. 5, *B* and *C*. The enhanced dynamics at the dimeric interface over the WT is clearly evident in DS1^R^-3^R^. In WT, the strong binding of C2 manifests as the absence of any significant motion. Amongst the reduced systems, the loss in structural symmetry as well as enhancement in cluster dynamics over WT is found to be lowest in DS4^R^.

**Figure 5.**
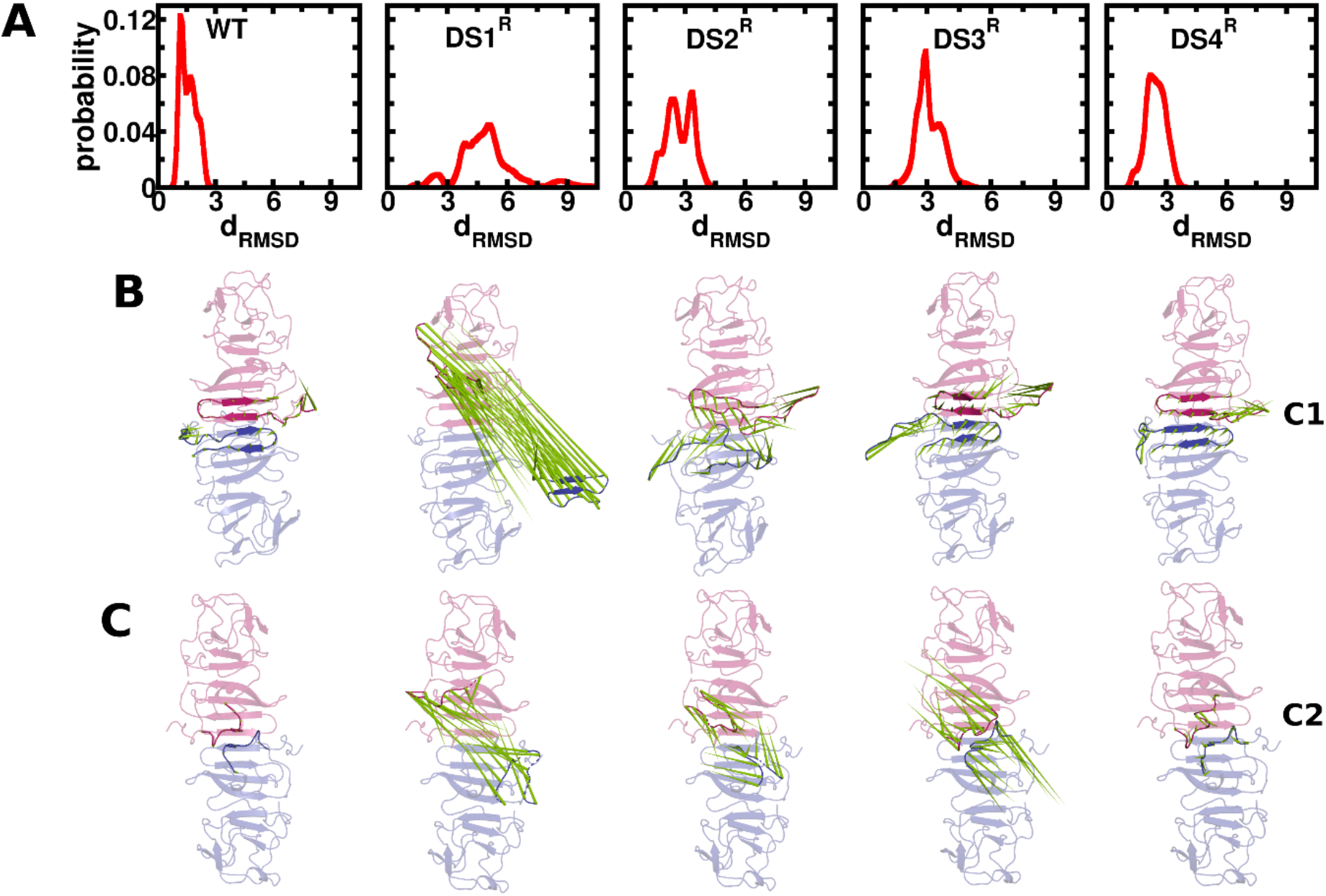
Evidence of structural asymmetry in the β-ladder homodimer. Normalized distribution of differential RMSD (d_RMSD_; in Å) (*A*); and corresponding representation of essential dynamics of C1 and C2 as a porcupine plot for WT and DS1^R^-4^R^ (*B* and *C*). In the porcupine plot, the cluster regions are superposed over the starting conformation.

As mentioned earlier, homooligomerization is associated with minimization of surface exposure to surrounding solvent (1). In Fig. 6 *A*, we compare distributions of the solvent accessible surface area (SASA) of the entire dimer in the WT and DS1^R^-4^R^. The increased solvent accessibility for the reduced systems emerge clearly, with more pronounced effect in the C2 cluster. We further examined the hydration numbers in the vicinity of common residues participating in C1 and C2. We find a noticeable increase in the hydration number (N_hyd_) for DS1^R^-3^R^, consistent with the observation of increased SASA in these systems. In Fig. 6 *B*, we have depicted the average hydration number with representative snapshots. Enhanced interfacial hydration is generally associated with stability loss in structural salt bridges (58). Therefore, we evaluated the symmetric persistence of the internal Asp^197^-Lys^221^ salt bridge in the vicinity of C2. As shown in Fig. S7, this salt bridge shows remarkable co-persistence in both monomers in the WT and DS4^R^ along the simulation trajectories. In DS1^R^-3^R^, however, there is a strong propensity for asymmetric loss of this salt bridge. As seen in Fig 4, the loss of salt bridge persistence in these systems reflects the asymmetric disruption in the C2 cluster that accompanies significantly enhanced hydration around the cluster residues with weakened contact.

**Figure 6.**
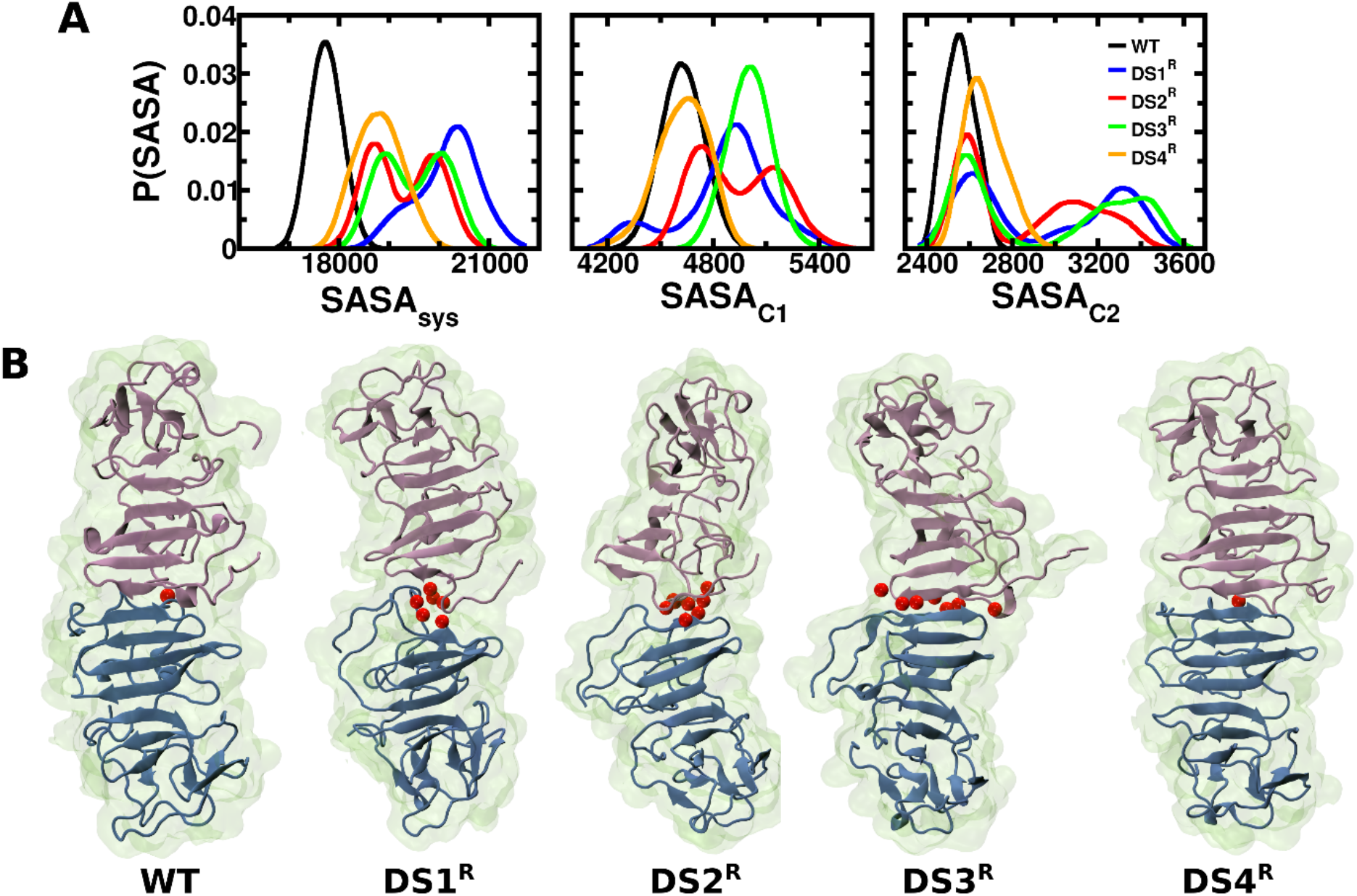
Solvent accessible surface area (SASA, in Å^2^) distributions for the entire dimeric system and the clusters (*A*), and representative conformation depicting the average number of interfacial water molecules in red (*B*).

### IV. Entropic signatures of homooligomeric asymmetry

In proteins, conformational entropy (S_conf_) is a key thermodynamic signature whose variation can signal structural rearrangement, altered stability, and altered interaction potential. Proteinprotein or protein-ligand binding is typically associated with a decrease in S_conf_ although there are known cases of accompanying entropic increase (59,60). The direction of S_conf_ change crucially contributes to the overall direction of enthalpy-entropy compensation.

Barring effects of thermal fluctuation, homooligomerization can be expected to result in high degrees of symmetry in S_conf_ of participating monomers. However, our earlier analyses showed that distal disulfide reductions trigger asymmetric dynamics within the NS1 β-ladder dimer. Herein, we report the entropic manifestations of the earlier observations. We point out that despite its importance, accurate estimation of S_conf_ remains a challenge with traditional methods (61). Herein, we leverage a highly accurate, model-free NMR based formalism based on the recognition that Sconf relates to the fast internal motions of a protein (40,62). The backbone (S^BB^) and sidechain (S^SC^) conformational entropies were calculated *via* the generalized NMR order parameter formalism (see Methods). First, considering the entire dimeric unit, we observe a net increase in both S^BB^ and S^SC^ in DS1^R^-4^R^ over the WT (see Table 2). We next evaluated the differences in backbone and sidechain entropy between the monomers along individual trajectories and their corresponding distributions; see Fig. 7 (for ΔS^BB^) and Fig. S8 (for ΔS^SC^). The data demonstrate enhanced entropic difference between the monomers during temporal evolution of DS1^R^-4^R^. The differences are further demonstrated from the nature of the distributions. The median values of ΔS^BB^ are closer to 0, whereas those for DS1^R^-4^R^ show a greater spread and enhanced skew. We note that the entropic asymmetry is relatively more pronounced in ΔS^BB^. However, though significantly smaller in magnitude (63), enhanced ΔS^SC^ asymmetry over WT is also evident in DS1^R^-4^R^.

**Table 2.** Change of backbone (ΔS^BB^) and sidechain (ΔS^SC^) entropies in cal mol^−1^ K^−1^. The subscripts describe entropic differences from WT (DS-WT); between monomers (M); between monomeric residues comprising the clusters C1 (C1), C2 (C2), and C2’ (C2’). See the text for details. Median and mean positions with standard deviations in braces are reported where relevant.

| $\Delta S_{\text{conf}}$ | | WT | DS1<sup>R</sup> | DS2<sup>R</sup> | DS3<sup>R</sup> | DS4<sup>R</sup> |
| --- | --- | --- | --- | --- | --- | --- |
| $\Delta S^{\text{BB}}_{\text{DS-WT}}$ | | - | 225.52 | 143.29 | 191.62 | 132.96 |
| $\Delta S^{\text{SC}}_{\text{DS-WT}}$ | | - | 0.6 | 0.24 | 0.41 | 0.12 |
| $\Delta S^{\text{BB}}_{\text{M}}$ | Median | 2.92 | -3.87 | -4.34 | -0.76 | -17.31 |
|  | Mean | 1.7 (14.5) | 7.2 (48.5) | -8.9 (33.4) | 2.79 (30.0) | 15.78 (29.1) |
| $\Delta S^{\text{SC}}_{\text{M}}$ | Median | 0.021 | 0.025 | 0.038 | -0.02 | 0.037 |
|  | Mean | -0.002 (0.25) | 0.01 (0.34) | 0.07 (0.31) | -0.03 (0.34) | 0.06 (0.33) |
| $\Delta S^{\text{BB}}_{\text{C1}}$ | Median | 0.5 | 4.4 | 6.00 | -0.1 | 5.1 |
|  | Mean | 0.6 (6.3) | 4.3 (10.4) | 5.6 (7.1) | -1.8 (9.2) | 5.1 (5.6) |
| $\Delta S^{\text{SC}}_{\text{C1}}$ | Median | -0.03 | -0.03 | -0.05 | -0.01 | 0.01 |
|  | Mean | -0.03 (0.1) | 0.02 (0.12) | 0.06 (0.12) | -0.01 (0.12) | -0.2 (0.1) |
| $\Delta S^{\text{BB}}_{\text{C2}}$ | Median | -0.15 | -2.05 | -3.04 | -0.87 | 1.91 |
|  | Mean | -0.4 (1.7) | 3.5 (12.5) | -1.5 (6.04) | -0.77 (6.07) | 1.61 (4.9) |
| $\Delta S^{\text{SC}}_{\text{C2}}$ | Median | -0.003 | -0.029 | -0.038 | -0.018 | 0.001 |
|  | Mean | 0.01 (0.1) | -0.02 (0.2) | -0.03 (0.2) | -0.04 (0.2) | -0.01 (0.2) |
| $\Delta S^{\text{BB}}_{\text{C2'}}$ | Median | - | 4.43 | -2.57 | -0.46 | 1.82 |
|  | Mean | - | 4.93 (3.6) | -1.54 (3.2) | -0.62 (2.6) | 1.97 (1.9) |
| $\Delta S^{\text{SC}}_{\text{C2'}}$ | Median | - | -2.11 | 1.0 | -3.2 | -1.06 |
|  | Mean | - | -2.1 (0.05) | 0.98 (0.11) | -3.2 (0.14) | -1.07 (0.11) |

**Figure 7.**
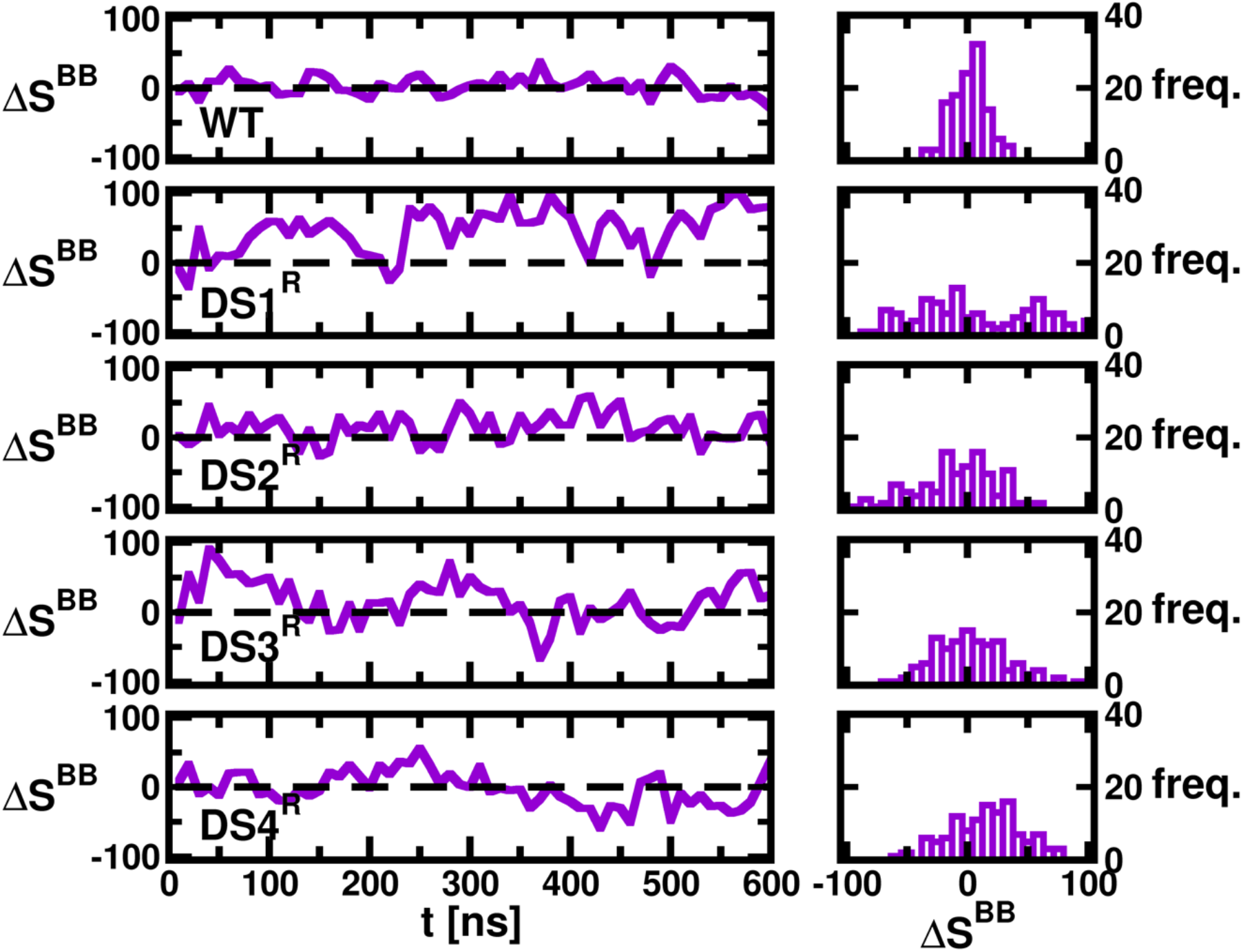
Temporal signature of differential backbone entropy (ΔS^BB^; in cal mol^−1^ K^−1^) between the monomers of β-ladder dimer of the five systems (*left panel*). Distributions of ΔS^BB^ for corresponding systems (*right panel*).

We next estimated the entropy differences (ΔS^BB^ and ΔS^SC^) between the monomers that comprise the clusters of the dimer interface. The analysis, presented in Fig. S9 and Fig. S10 for C1, clearly show an increased entropic asymmetry between the monomers in the disulfide reduced systems over the WT. As earlier, the differences are more pronounced in ΔS^BB^ compared to ΔS^SC^. We note that the median value of ΔS^BB^ is narrowly clustered around 0 for the WT; greater spread is noticeable in DS1^R^-4^R^. We have observed previously that the electrostatically stabilized C2 is the predominant contributor to interfacial symmetry and stability in the WT dimer, and further, is highly susceptible to allosteric perturbation and distortion triggered with elimination of internal disulfides. In Fig. 8, we have compared the temporal evolution and the distributions of ΔS^BB^ of residues comprising the WT C2 cluster for the simulated systems. Noting that allosteric distortion in C2 leads to the formation of new contacts (see Fig 4), we have additionally evaluated ΔS^BB^ between the monomers for the distorted interface referred to as C2’. The corresponding plots for ΔS^SC^ are presented in Fig. S11. We observe that entropic differences between the monomers in WT are markedly close to 0 and has a very low spread in the distributions. This symmetry is clearly lost for the reduced systems which exhibit a marked spread and enhanced skew in both ΔS^BB^ and ΔS^SC^. For C2, although magnitudes of ΔS^BB^ are higher and the asymmetry clearly pronounced in ΔS^BB^, the asymmetry is evident in ΔS^SC^ as well. Interestingly, when C2’ is considered, the median position in ΔS^SC^ shifts noticeably from 0 for all disulfide reduced systems; this shift is most prominent for DS3^R^. The median, mean, and corresponding standard deviations of all ΔS_conf_ calculations are provided in Table 2.

**Figure 8.**
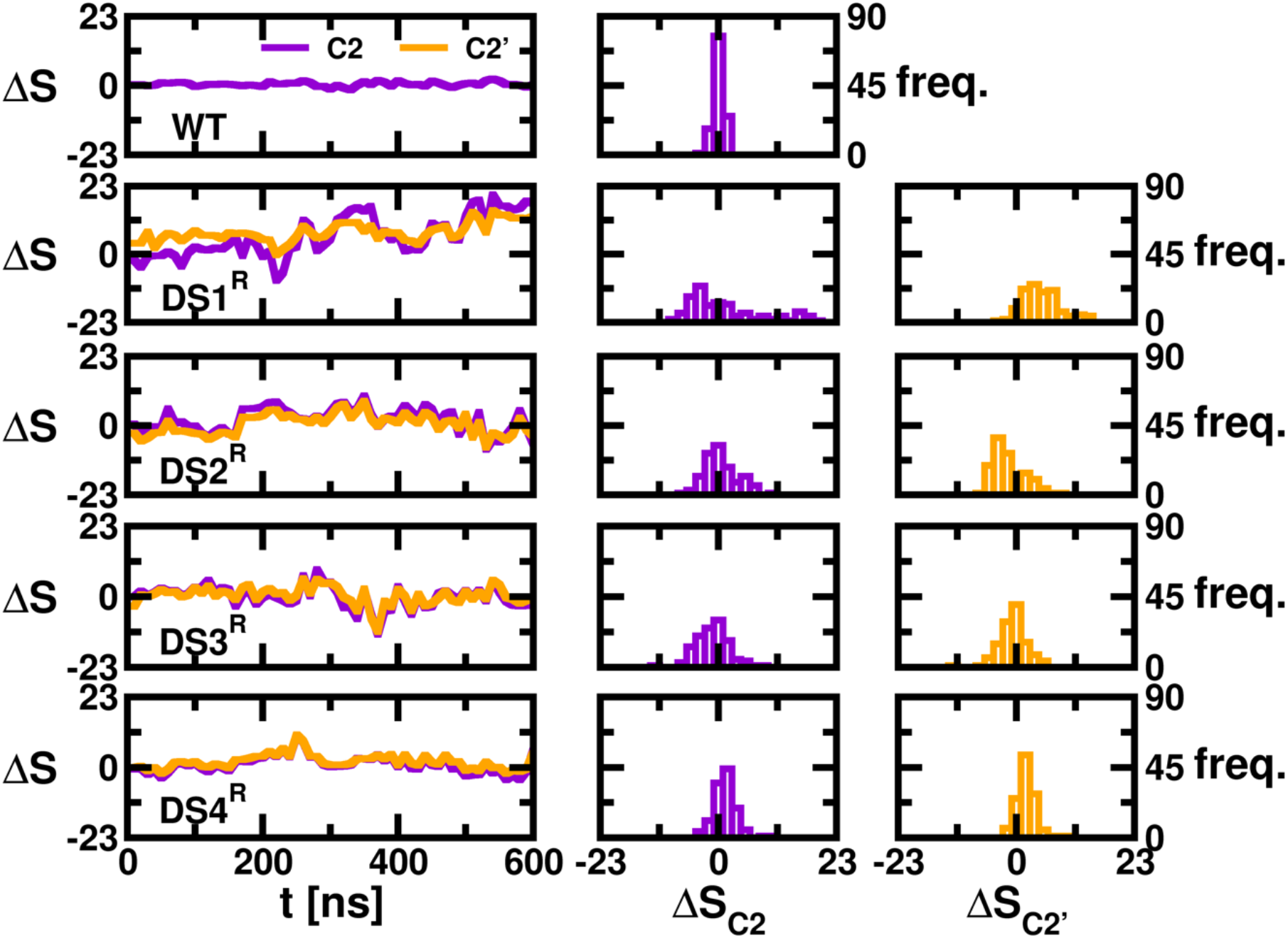
Time evolution of differential backbone entropy for native contacts of C2 (Color: Purple) of each system and the altered contacts (i.e. C2’; Color: orange) of reduced systems. Contacts with an average strength exceeding −2 kcal mol^−1^ are considered (*left panel*). Distributions of ΔS_C2_ (*middle panel*) and ΔS_C2’_ for corresponding systems (*right panel*). Units in cal mol^−1^K^−1^

The distinct entropic asymmetry triggered allosterically by the disulfide reductions commensurate with asymmetric structural dynamics can be expected to attribute differential binding affinity of the individual NS1 monomers. This should result in greater structural plasticity and heterogeneity, and manifest in low symmetry in the higher ordered oligomeric states. It is further likely that the entropic asymmetry will lead to differences in binding potentials of interfacial residues with candidate drug molecules; this effect may be leveraged to amplify the asymmetric effect and further hinder higher ordered homooligomerization. Harnessing these effects cumulatively could lead to significant disruptions in NS1 oligomerization pathways that are necessitated in viral replication and pathogenesis.

## Conclusion

Viral genomes are typically small and not amenable to the translation of large, multi-domain proteins, and several crucial viral processes rely on the homooligomerization of smaller protein units. In ZIKV, higher ordered homooligomerization (trimerization of dimers) of the non-structural NS1 protein relates to pathogenesis, and thereby necessitates determination of interventional schemes to its self-assembly. Motivated thus, herein we have presented analyses of conformational ensembles generated *via* extensive molecular simulations that provide evidence of allosteric destabilization of the dimeric form of the primary β-ladder domain of the NS1 protein. The dimeric form is stabilized principally by two non-covalent interfacial clusters, each of which is predominantly stabilized either hydrophobically or electrostatically. Subjecting each monomeric unit to a relatively small, symmetric perturbation, namely disulfide reduction in one of the four possible location pairs, allosterically triggers structural and thermodynamic destability at the binding interfaces. Importantly, our analyses reveal that the perturbation triggered symmetrically is commensurate with large variance in properties of the two monomeric units. The symmetry in conformation and essential dynamics observed in the WT is clearly broken in most of the disulfide reduced systems. The asymmetry triggered by the allosteric perturbation also manifests in the asymmetric distribution and wider spread of the backbone and sidechain conformational entropies of the two monomeric units.

The implications of this work are manifold and should lead to further *in vivo, ex vivo*, and *in silico* studies towards the design of antiviral strategies. First, while the β-ladder domain constitutes the major binding domain of the NS1 assembly, the effects on its weakening on the full dimeric organization should be probed in detail. Secondly, the structural and thermodynamic aspects of hexameric organization of the unperturbed, as well as the distorted dimers within lipid bilayers, must be mechanistically explored. Furthermore, in light of our discovery of intra-monomeric disulfide reduction distorting the dimeric assembly, it would be useful to investigate the effects of pH alterations on the organizational stability of the NS1 protein and its effects downstream on ZIKV virulence. It is possible, however, that the allosteric networks underlying NS1 organization are susceptible to more direct molecular interference, specifically *via* drug binding. Exploiting the potential for allosteric disruption, entropic redistribution and loss of symmetry in conjunction with differential drug binding affinities of individual monomers could lead to novel ways of weakening the stabilizing interaction networks. Some of these investigative efforts are underway in our group. Prior to the conclusion, we point out that in recent years, novel and highly infectious viral diseases have frequently gained pandemic proportions. It may be worthwhile considering allosteric interventions that exploit a potential weakness of the viral genome, namely its frequent reliance on protein oligomerization for unleashing virulence.

## Author contributions

P.R. and N.S. designed the research, analyzed the data, and wrote the manuscript. P.R. did the simulation. S.R. contributed in the code writing.

## Acknowledgments

P.R. acknowledges CSIR for her Junior Research Fellowship. SR acknowledges DST for his INSPIRE fellowship. N.S. acknowledges Science and Engineering Research Board (SERB) for the funds used to procure computational resources (EMR/2016/001108). Additional resources provided by PARAM Yuva II-CDAC supercomputing facility in Pune is acknowledged.

